# Sympathetic NPY controls glucose homeostasis, cold tolerance, and cardiovascular functions in mice

**DOI:** 10.1101/2023.07.24.550381

**Authors:** Raniki Kumari, Raluca Pascalau, Hui Wang, Sheetal Bajpayi, Maria Yurgel, Kwaku Quansah, Samer Hattar, Emmanouil Tampakakis, Rejji Kuruvilla

**Affiliations:** Department of Biology, Johns Hopkins University, Baltimore, Maryland, 21218, USA; Section on Light and Circadian Rhythms, National Institute of Mental Health, National Institutes of Health, Bethesda, Maryland, 20892, USA; Division of Cardiology, Johns Hopkins School of Medicine, Baltimore, Maryland, 21205, USA

## Abstract

Neuropeptide Y (NPY) is best known for its effects in the brain as an orexigenic and anxiolytic agent and in reducing energy expenditure. NPY is also co-expressed with Norepinephrine (NE) in sympathetic neurons. Although NPY is generally considered to modulate noradrenergic responses, its specific roles in autonomic physiology remain under-appreciated. Here, we show that sympathetic-derived NPY is essential for metabolic and cardiovascular regulation in mice. NPY and NE are co-expressed in 90% of prevertebral sympathetic neurons and only 43% of paravertebral neurons. NPY-expressing neurons primarily innervate blood vessels in peripheral organs. Sympathetic-specific deletion of NPY elicits pronounced metabolic and cardiovascular defects in mice, including reductions in insulin secretion, glucose tolerance, cold tolerance, pupil size, and an elevation in heart rate, while notably, however, basal blood pressure was unchanged. These findings provide new knowledge about target tissue-specific functions of NPY derived from sympathetic neurons and imply its potential involvement in metabolic and cardiovascular diseases.

## Introduction

The sympathetic nervous system is a major conduit for communication between the brain and the periphery. While widely known for its role in triggering “fight or flight responses” in the event of danger or stress, the sympathetic nervous system is also vital for maintaining body homeostasis during daily activities such as feeding or exercising. Organized into a system of ganglia and nerves in the peripheral nervous system (PNS), sympathetic neuron cell bodies, resident in ganglia, send long axonal projections throughout the body to innervate diverse peripheral organs and tissues to control a broad spectrum of physiological processes including heart rate, blood pressure, body temperature, blood glucose levels, and immune function (Scott-Solomon et al., 2021). Homeostatic and adaptive sympathetic regulation of different peripheral organs is remarkably precise, suggesting the existence of dedicated neural circuits that drive different effector organ functions (Janig and Habler, 2000). However, there is limited understanding of the functional diversity of sympathetic neurons at the molecular and cellular level.

It has been proposed that one basis for the functional diversity of sympathetic neurons is the combinatorial expression of neuropeptides, along with a classical neurotransmitter, which provides a “neurochemical code” that allows for the innervation of distinct target organs and the regulation of specific autonomic functions (Ernsberger et al., 2020). Neuropeptide Y (NPY) is a 36-amino acid peptide that is co-stored and co-released with norepinephrine (NE) from sympathetic neurons and adrenal glands (Ekblad et al., 1984; Lundberg et al., 1989a; Lundberg et al., 1982; Whim, 2006). In contrast to the vast amount of knowledge of the potent effects of NPY expressed in brain regions, specifically hypothalamic and brainstem nuclei in regulating food intake (Kalra and Kalra, 2004; Zarjevski et al., 1993), mood (Alldredge, 2010), energy balance (Loh et al., 2015), and cardiovascular functions (Tan et al., 2018; Walker et al., 1991), the actions of NPY in peripheral sympathetic neurons have been relatively under-studied. NPY functions have been largely studied at the whole-organism level using global knockout mice (Erickson et al., 1996a; Erickson et al., 1996b; Karl et al., 2008; Lin et al., 2004; Loh *et al*., 2015) or by pharmacological manipulations (Zhang et al., 2011), and the knowledge of specific roles for NPY in defined neuronal populations remains incomplete.

In sympathetic neurons, electrophysiological and pharmacological studies on tissue preparations indicate that NPY is co-released with NE from nerve terminals, specifically during prolonged or high-frequency stimulation (Lundberg *et al*., 1989a). Sympathetic-derived NPY has mild vasoconstrictor effects on its own (Ekblad *et al*., 1984; Lundberg *et al*., 1982) although it powerfully potentiates NE-induced vascular contractile responses by acting post-synaptically (Dahlof et al., 1985; Ekblad *et al*., 1984; Hakanson et al., 1986; Lundberg et al., 1985; Wahlestedt et al., 1985). NPY, released from sympathetic nerves, also exerts a pre-synaptic effect in inhibiting NE release (Lundberg *et al*., 1985; Wahlestedt and Hakanson, 1986). More recent studies with genetic ablation of NPY and NE co-expressing sympathetic neurons suggest roles for these neurons in modulating inflammatory responses (Yu et al., 2022) and cardiac excitability (Sharma et al., 2023) in mice. However, whether observed phenotypes were due to the loss of NPY, or NE, or other neuron-derived factors could not be delineated from these studies. Thus, other than its classical role as a modulator of sympathetic neurotransmission, the specific functions of NPY expressed in sympathetic neurons remain to be elucidated.

Here, using an intersectional genetic labeling approach, we show that sympathetic neurons co-expressing NPY and NE are more abundant in prevertebral, compared to paravertebral, sympathetic ganglia. Axons of NPY and NE co-expressing sympathetic neurons extend along, and primarily innervate, blood vessels in peripheral tissues. Using conditional knockout mice, we show that loss of NPY from sympathetic neurons does not affect neuron development, target innervation, or activity, but results in pronounced metabolic and cardiovascular defects, including reductions in insulin secretion, glucose tolerance, cold tolerance, pupil size, and an elevation in heart rate. Surprisingly, despite the well-documented role of NPY in vasoconstriction, we found that blood pressure under basal conditions was unaffected in mutant mice. We also found elevated circulating NE levels in NPY conditional knockout mice, which may be due, in part, to increased NE biosynthesis in adrenal glands. Together, these findings provide fresh insight into specific actions of NPY in the sympathetic nervous system.

## Results

### NPY is expressed in a higher proportion of sympathetic neurons in prevertebral ganglia

Anatomically, the sympathetic nervous system is organized into paired paravertebral ganglia located in chains along the rostro-caudal axis on both sides of the spinal cord, and single prevertebral ganglia located at the midline, ventral to the spinal cord. Although NPY is widely considered to be a co-transmitter with NE in sympathetic neurons (Hakanson *et al*., 1986), previous studies have suggested that NPY expression is not uniform among noradrenergic sympathetic neuron populations (Furlan et al., 2016; Liu et al., 2020; Masliukov et al., 2012). Here, we used a genetic labeling approach to assess the distribution of NPY-expressing sympathetic neurons. *NPY^Cre^* mice (Milstein et al., 2015) were crossed with *ROSA26^eYFP^* mice (Srinivas et al., 2001) in conjunction with immunostaining for Tyrosine Hydroxylase (TH) to mark noradrenergic neurons. We chose the Superior Cervical Ganglia (SCG), the rostral-most ganglia in the sympathetic chain, and the Celiac–Superior Mesenteric Ganglia (CG-SMG), which lie within the abdominal cavity, to represent para- and pre-vertebral ganglia, respectively. SCG neurons project to the head and neck vasculature, eye, pineal glands, salivary glands and skin overlying the head and neck, while CG-SMG neurons project to abdominal organs including liver, pancreas, spleen, kidneys, stomach, and the proximal small intestine.

We found that ∼90% of TH-positive sympathetic neurons expressed the EYFP reporter in CG-SMG (**Figures 1B, F**), compared to only ∼43% in the SCG in postnatal day 21 old mice (**Figures 1A, E**). Since the genetic reporter labels all cells that expressed *Npy* at any moment during development, including any that may have subsequently down-regulated NPY protein expression, we also performed NPY immunostaining. Similar to observations with genetic labeling, the majority of TH-positive neurons showed NPY immunoreactivity in the CG-SMG, compared to partial overlap in the SCG at postnatal day 21 (**Figures 1C, D**). Quantification revealed that 25% of TH-positive neurons were GFP^+^;NPY^-^ in the SCG compared to 6.3% in the CG-SMG, suggesting that NPY protein expression is down-regulated in a significant proportion of sympathetic neurons in the SCG but not in CG-SMG.

**Figure 1.**
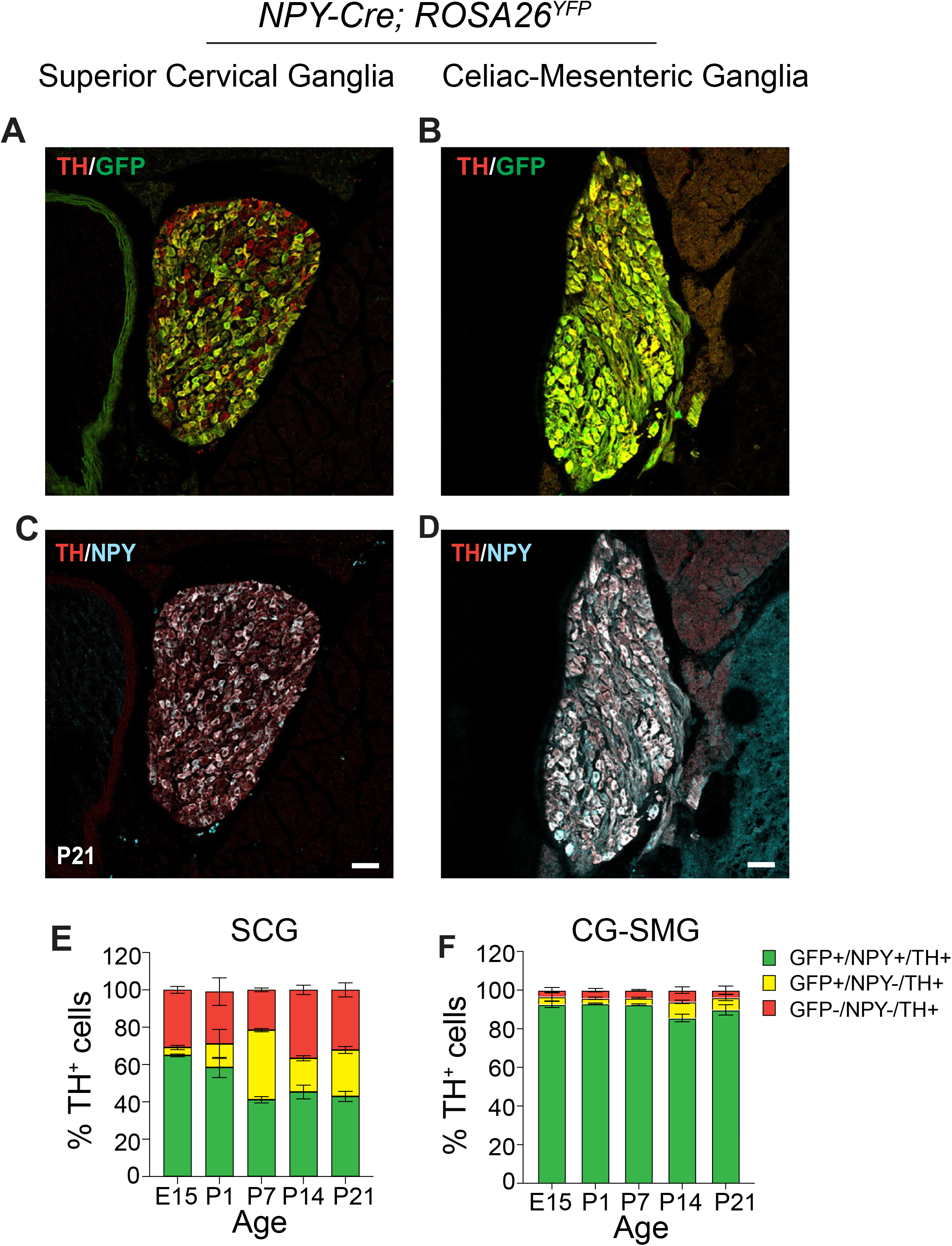
Differential NPY expression in paravertebral and prevertebral sympathetic neurons. **(A-D)** Immunostaining for TH, NPY, and GFP in Superior Cervical Ganglia (SCG, paravertebral) (**A, C**) and Celiac-Superior Mesenteric Ganglia (CG-SMG, prevertebral) (B, D) in NPY-Cre:*ROSA26^eYFP^* reporter mice. **(E, F)** Percentage of NPY-expressing noradrenergic neurons in SCG (E) and CG-SMG (F). NPY and TH are co-expressed (green bar) in ∼65% of SCG neurons at E15 and downregulated to ∼43% at P21. NPY and TH are co-expressed in ∼90% of CG-SMG neurons throughout development. Neurons that expressed NPY earlier, but no longer do so at the time of examination, are represented by the yellow bars. Sympathetic neurons that never express NPY are indicated by red bars. n=3 mice per genotype for each time-point. Scale bars, 50 μm.

To examine changes in the proportion of NPY-expressing neurons in sympathetic ganglia over time, we performed genetic and immunofluorescence labeling at various developmental time-points in mice. We found that the percentage of NPY-immunoreactive (GFP^+^;NPY^+^) noradrenergic neurons (∼90%) remains constant across all developmental time points examined (embryonic day 15, postnatal days 1, 7, 14, and P21) in the CG-SMG (**Figure 1F**). In contrast, in the SCG, the percentage of NPY-expressing sympathetic neurons decreases from 65% at E15 to 41% at P7, and then stays constant at later postnatal stages examined (P14 and P21) (**Figure 1E**). The decrease in the NPY-immunoreactive population in the SCG is likely due to down-regulation of NPY expression and not cell loss, because the percentage of genetically labeled neurons (GFP-positive) remained largely unchanged over time (**Figure 1E**). Further, of note, almost one-third (32%) of TH-positive neurons in the SCG were neither NPY nor GFP-positive when examined at P21, while this population was minimal (3.6%) in the CG-SMG (**Figures 1E, F**).

Together, these results indicate that the proportion of NPY-expressing noradrenergic neurons is markedly higher in pre-*versus* para-vertebral sympathetic ganglia. Further, in prevertebral neurons, the adult pattern of NPY and NE co-expressing neurons is acquired early at embryonic stages and maintained throughout development, compared to paravertebral neurons where the mature pattern of NPY expression is only established after birth.

### NPY-expressing sympathetic axons largely innervate blood vessels

Given the differential abundance of NPY-expressing neurons in para- and pre-vertebral sympathetic ganglia, we next wanted to define their axonal projections to diverse peripheral organs and tissues. NPY is expressed in many non-neuronal cells in peripheral tissues, including blood platelets (Myers et al., 1988), white adipose tissue (Yang et al., 2008), bone cells (Baldock et al., 2007), and pancreatic beta-cells (Rodnoi et al., 2017), which would have been labeled by the *NPY^Cre^*;*ROSA26^eYFP^* reporter mice or NPY immunostaining (**see Figure S1A**) making it difficult to distinguish the signal coming from innervating axons. To specifically visualize NPY-expressing sympathetic nerves, we used an inter-sectional genetic approach. We crossed a dual recombinase-dependent reporter mouse line, *Rosa26^LSL-FSF-tdTomato^ (Ai65)* mice (Madisen et al., 2015), to *DBH^Cre^* (Quach et al., 2013) and *NPY^FLP^* mice (Daigle et al., 2018) (**Figure 2A**). In *DBH^Cre^*;*NPY^FLP^*;*ROSA26^LSL-FSF-tdTomato^*mice (henceforth referred to as *NPY^DBH^*-*TdTomato* mice), tdTomato reporter expression is restricted to NPY^DBH^ sympathetic neurons co-expressing NPY and Dopamine-beta-hydroxylase (DBH), an enzyme in the NE biosynthetic pathway.

**Figure 2.**
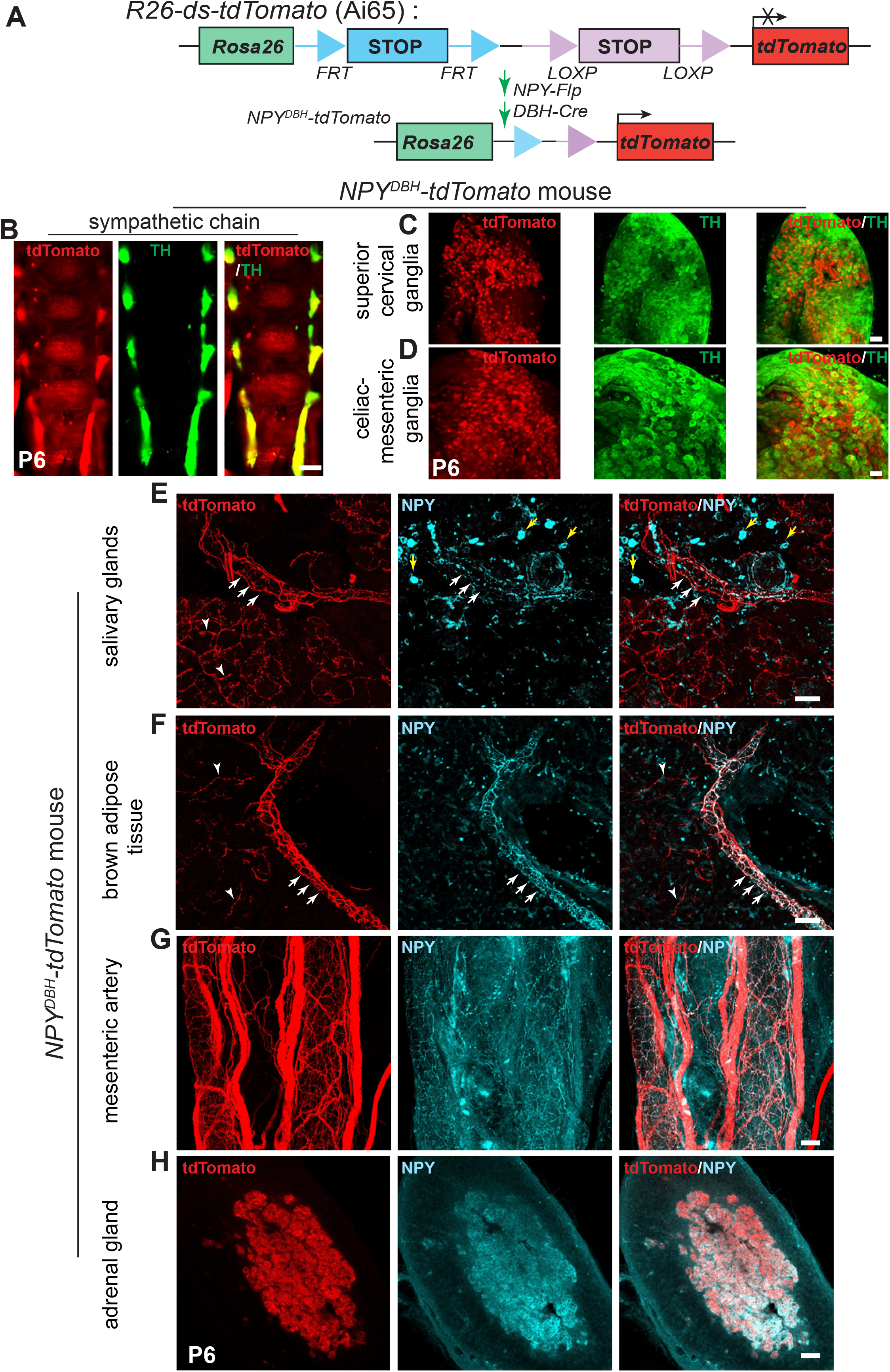
NPY-expressing sympathetic axons primarily innervate blood vessels. **(A)** Schematic showing the intersectional genetic strategy to generate *NPY^DBH^-tdTomato* mice, where tdTomato expression is activated by removal of two *STOP* cassettes by Flpo and Cre recombinases in sympathetic neurons co-expressing NPY and DBH. **(B-D)** Co-localization of tdTomato and TH in sympathetic chain (B), SCG (C) or CG-SMG (D) using whole-mount immunostaining in *NPY^DBH^-tdTomato* mice at post-natal day 6. Scale bar, 500 μm (B) and 100 μm (C, D). **(E, F)** Innervation of paravertebral targets (salivary glands, BAT) by NPY^DBH^-tdTomato sympathetic fibers. TdTomato signal (red) is observed in axons aligned with blood vessels (white arrows) as well as in tissue parenchyma (white arrowheads). Compared to reporter expression, axonal NPY immunostaining (cyan) is only found in axons associated with vasculature. Yellow arrows indicate NPY immunostaining in non-neuronal cells. Scale bar, 50 μm. **(G, H)** Co-localization of NPY protein and tdTomato reporter expression in mesenteric artery (prevertebral sympathetic target, G) and adrenal chromaffin cells (H). Scale bar, 50 μm.

We observed NPY reporter expression in paravertebral sympathetic chain ganglia distributed along the rostro-caudal axis in postnatal (P6) mice as visualized by whole-mount TH and Red Fluorescent Protein (RFP) immunostaining and light sheet microscopy (**Figure 2B**). Similar to the observations in *NPY^Cre^*;*ROSA26^eYFP^* mice, reporter expression showed partial overlap with TH immunoreactivity in para-vertebral ganglia (SCG), and almost complete overlap with TH in pre-vertebral ganglia (CG-SMG) in *NPY^DBH^*-*TdTomato* mice (**Figure 2C, D**).

To visualize innervation of peripheral organs by NPY-expressing sympathetic axons, we performed immunostaining using antibodies against tdTomato, TH, or NPY in tissue sections or whole-mounts from *NPY^DBH^*-*TdTomato* mice at postnatal day 6. In salivary glands and interscapular brown adipose tissue (BAT), which are innervated by paravertebral ganglia neurons in the SCG and stellate/thoracic chain ganglia, respectively, we observed tdTomato reporter expression in axons extending along blood vessels and also in finer branches within the tissue parenchyma (**Figures 2E, F**). However, compared to reporter expression, NPY immunoreactivity was only present in axons associated with blood vessels (**Figures 2E, F**). These results suggest that the developmental down-regulation of NPY expression specifically occurs in neurons that innervate the tissue parenchyma, and that vascular-derived signals may be required for the maintenance of NPY expression in sympathetic neurons. TH immunostaining showed partial co-localization of TH immunoreactivity with reporter signal in axons innervating salivary glands and brown adipose tissue (**Figures S1B, C**), consistent with the pattern of co-expression of TH and NPY in neuronal cell bodies in paravertebral ganglia at postnatal stages.

For prevertebral targets, we assessed innervation of the mesenteric artery, a major artery supplying the gastro-intestinal tract, as well as the pancreas, which receives innervation from CG-SMG neurons (Ahren, 2000). TdTomato labeling showed a dense meshwork of NPY-expressing fibers around the mesenteric artery and aligned along smaller blood vessels in the pancreas (**Figures 2G and Figure S1A**). NPY protein expression largely overlaps with tdTomato reporter signal (**Figures 2G and Figure S1A**), as well as with TH immunoreactivity (**Figures S1D, E**), consistent with the observations that the majority of pre-vertebral ganglia neurons co-express NPY and TH, and that NPY protein expression is not down-regulated during development as in the para-vertebral ganglia.

In addition to sympathetic neurons, NPY is also expressed in adrenal chromaffin cells (Lundberg et al., 1983; Renshaw and Hinson, 2001; Wolfensberger et al., 1995), which are neuroendocrine cells that primarily secrete epinephrine, but also norepinephrine, to regulate myriad cardiovascular and metabolic processes, particularly in response to stressful stimuli. Similar to pre-vertebral neurons, we also observed a complete overlap between tdTomato reporter expression, NPY, and TH immunostaining in adrenal chromaffin cells in *NPY^DBH^-TdTomato* mice at postnatal day 6 (**Figure 2H and Figure S1F**). These results suggest that the developmental regulation of NPY expression in adrenal glands resembles the scenario in pre-vertebral ganglia neurons.

Together, these results suggest that NPY-expressing sympathetic neurons largely innervate blood vessels in peripheral target tissues innervated by both pre- and para-vertebral ganglia neurons. However, the majority of axons from prevertebral noradrenergic neurons maintain NPY expression during postnatal life compared to paravertebral neurons, consistent with the patterns observed in neuronal cell bodies.

### Loss of sympathetic NPY elicits soma atrophy but does not affect neuron survival or target innervation

To address the function(s) of NPY in sympathetic neurons, we generated conditional knockout mice, where mice with a floxed *NPY* allele (*NPY^fl/fl^* mice) (Wee et al., 2019) were crossed with *DBH^Cre^* mice. *DBH^Cre^;NPY*^fl/fl^ mice (henceforth referred to as NPY cKO mice) survived to adulthood, had no gross morphological abnormalities, and had normal body weight (see **Figure 5A**). Litter-mate *NPY*^fl/fl^ mice were used as controls in all analyses. NPY expression, visualized by immunofluorescence, was significantly reduced in mutant sympathetic (CG-SMG) ganglia (**Figure 3A**), and qPCR analysis indicated an almost complete loss of *Npy* transcript in para-vertebral (SCG) and pre-vertebral (CG-SMG) ganglia (**Figure 3B**). Given that adrenal chromaffin cells also co-express DBH and NPY (Wolfensberger *et al*., 1995), we saw a drastic reduction in *Npy* mRNA levels in adrenal glands in mutant mice (**Figure 3B**).

**Figure 3.**
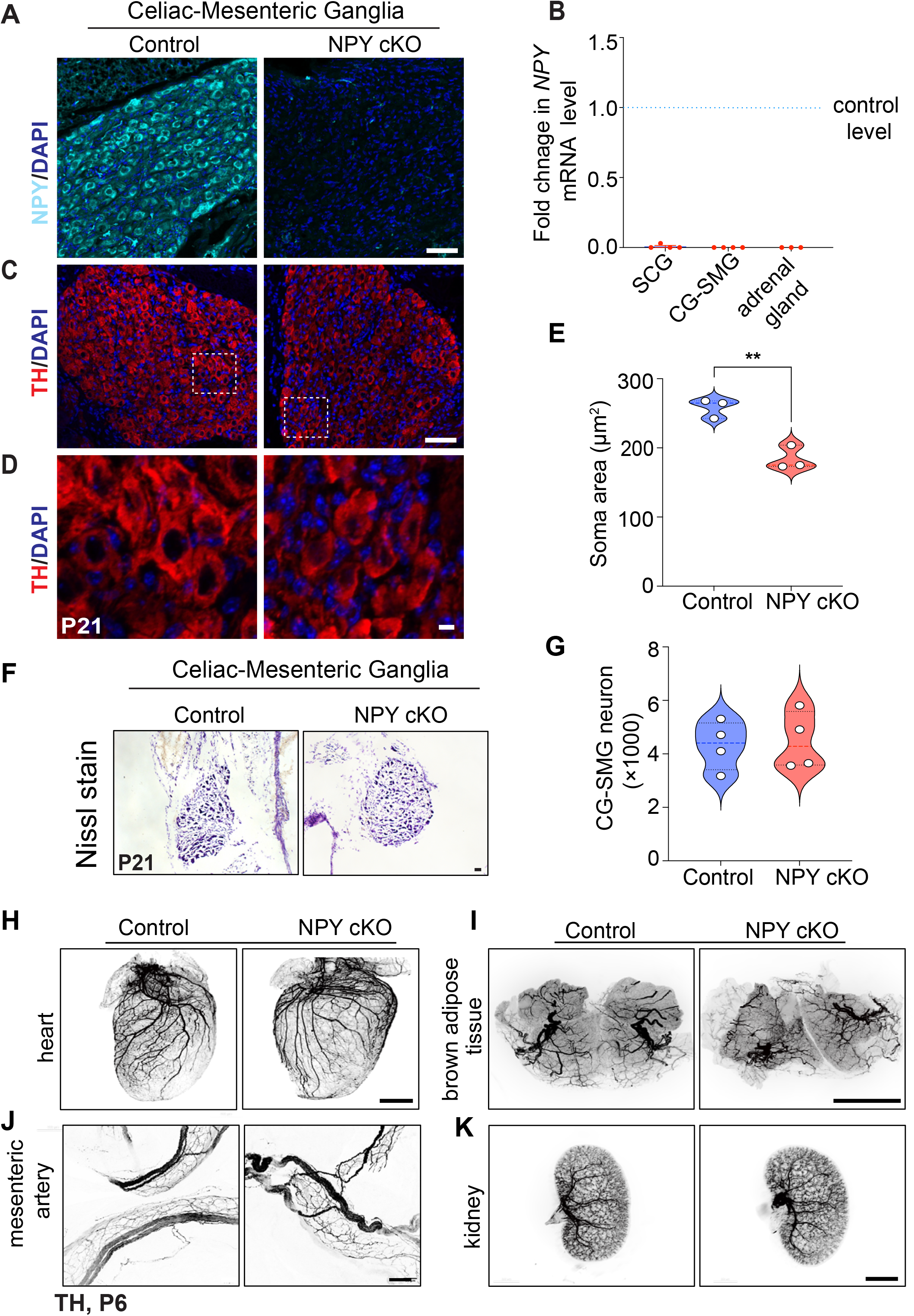
Loss of NPY in sympathetic neurons does not affect neuron survival and target innervation but results in reduced soma size. **(A)** Immunostaining shows a marked reduction in NPY protein in CG-SMG from NPY cKO mice compared to control mice. Scale bar, 50 μm. **(B)** qPCR analyses show a loss of *Npy* transcript in SCG, CG-SMG, and adrenal glands from NPY cKO mice. Results are means ± s.e.m from n= 4 animals per genotype for SCG, CG-SMG and n=3 animals per genotype for adrenal glands. (**C, D**) TH expression appears unaffected by NPY loss as shown by immunofluorescence. Shown in D are higher magnification views of the insets in C. Scale bar, 50 μm for C and 5 μm for D. **(E)** Quantification of soma areas from CG-SMG tissue sections shows that neuronal soma size is reduced in NPY cKO sympathetic ganglia. Results are means ± s.e.m from n=3 mice per genotype. **p<0.01, unpaired t-test. **(F)** Nissl staining of CG-SMG in control and NPY cKO mice. Scale bar, 50 μm. (**G**) Quantification of Nissl-stained neurons shows that neuron numbers are comparable between control and mutant mice. Results are means ± s.e.m from n=4 per genotype. **(H-K)** Whole organ TH immunostaining imaged by light sheet microscopy shows that sympathetic axon innervation of target tissues including the heart, brown adipose tissue, mesenteric arteries, and kidneys is comparable between NPY cKO and control mice. Scale bars, mesenteric arteries-50 μm; heart, and kidney-800 μm, brown adipose tissue-1500 μm.

To ask if loss of NPY in sympathetic neurons affects their development, we visualized neuron morphology using TH immunostaining and quantified neuron numbers in sympathetic ganglia using Nissl staining in mice at P21. Loss of NPY had no effect on TH expression in sympathetic neurons (**Figure 3C)**, but elicited a marked atrophy of neuronal soma (**Figures 3D, E**). Soma areas from tissue sections were 258 ± 8 μm^2^ for control neurons *versus* 184 ± 10 μm^2^ for mutant neurons. Despite the soma atrophy, sympathetic neuron numbers were not significantly different from that in control litter-mates (4324 ± 455 in control *vs.* 4484 ± 539.5 in mutant mice) based on quantification of cell counts (**Figures 3F, G**). Given the higher proportion of NPY and NE-co-expressing neurons in the CG-SMG, we largely focused on these ganglia, although similar results were obtained in the SCG. Whole-organ TH immunostaining in iDISCO-cleared tissues and light sheet microscopy showed that axon innervation density in diverse target tissues including the heart, brown adipose tissue (paravertebral targets) as well as mesenteric arteries and kidney (prevertebral targets) were unaffected in NPY cKO mice (**Figures 3H-K**). Together, these results indicate that NPY is required for regulating soma size in sympathetic neurons but is dispensable for neuron survival and target innervation during development.

### NPY cKO mice have elevated circulating NE levels

Previous studies have documented that central NPY signaling suppresses sympathetic tone (Baldock et al., 2014; Shi et al., 2013; Shi et al., 2017), while NPY, derived from peripheral sources, exerts post- and pre-synaptic effects in potentiating NE-induced responses and inhibiting NE release, respectively (Lundberg *et al*., 1985; Westfall et al., 1987; Zukowska-Grojec, 1995). To address how the selective loss of NPY in sympathetic neurons affects neuron activity, we performed immunostaining for c-Fos, an immediate early transcription factor that serves as a reporter of neuronal activity (Sheng and Greenberg, 1990), in CG-SMG tissue sections. We found similar numbers of c-Fos-positive sympathetic neurons in control and NPY cKO mice (11.2 ± 1.0 in control *versus* 9.33 ± 1.3 in mutant) at room temperature (**Figures 4A, C, E**). Activation of the sympathetic nervous system by a brief (1 hr) cold exposure at 4°C significantly increased the number of c-Fos-positive cells by ∼4.7-6.2-fold in both control and mutant ganglia (53.0± 2.6 in control *versus* 57.6 ± 4.0 in mutant) (**Figures 4B, D, and E**). These results suggest that basal and cold-induced activity in sympathetic ganglia is not affected by the loss of NPY.

**Figure 4.**
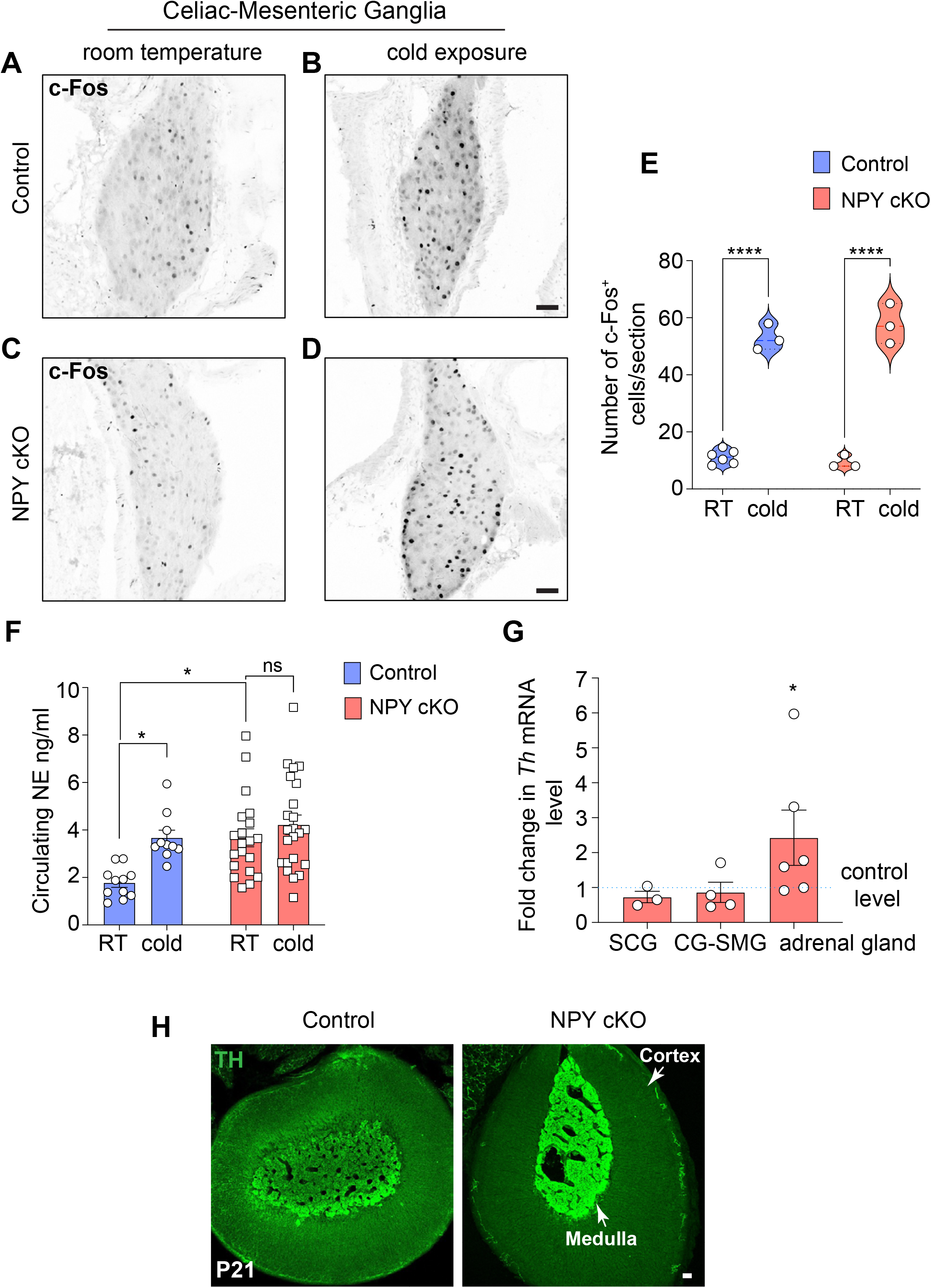
Loss of sympathetic NPY enhances circulating NE without affecting neuron activity. **(A-D)** Similar numbers of c-FOS-positive sympathetic neurons in CG-SMG from 6-8 week-old control and NPY cKO mice housed at room temperature (RT, 26°C). Cold exposure (4°C, 1 hr) increases the number of c-FOS-positive sympathetic neurons in both control and mutant ganglia. Scale bars, 50 μm. **(E)** Quantification of c-FOS-positive sympathetic neurons in control and mutant CG-SMG at RT and in response to cold exposure (4°C, 1 hr). Data are presented as means ± s.e.m from n=6 animals for control mice at RT, and n=3 animals for all other conditions. ****p p<0.0001; two-way ANOVA, Tukey’s multiple comparison test. **(F)** Plasma NE levels are significantly elevated in NPY cKO mice compared to control animals at room temperature (26°C). Cold exposure (4°C, 2 hr) significantly increases circulating NE in control, but not NPY cKO, animals. Data are as means ± s.e.m n=11 control and 20 mutant mice. *p<0.05; two-way ANOVA, Tukey’s multiple comparisons. **(G)** *Th* mRNA is increased in adrenal glands, but not in sympathetic ganglia, in NPY cKO mice relative to control animals as shown by qPCR analysis. Results are means ± s.e.m from n=3 mice per genotype for SCG, 4 for CG-SMG, and 6 for the adrenal glands. *p<0.05; one sample t-test. **(H)** TH immunofluorescence is increased in adrenal chromaffin cells in NPY cKO mice relative to control animals, indicating an increase in TH protein level. Scale bar, 50 μm.

To address the effects of NPY loss in sympathetic neurons on NE release, we measured circulating NE levels in NPY cKO and control mice kept at room temperature or exposed to 4°C for 2 hr. Plasma NE levels are thought to be primarily derived from sympathetic nerves, although a smaller contribution comes from secretion from adrenal chromaffin cells (Kvetnansky et al., 1979). At room temperature, we observed a significant increase (2.2-fold) in circulating NE in mutant mice compared to control animals (**Figure 4F**). In control animals, cold exposure at 4°C significantly increased plasma NE compared to room temperature (**Figure 4F**), consistent with elevated sympathetic activity. However, in NPY cKO animals, there was no further elevation in plasma NE in response to cold exposure (**Figure 4F**).

In addition to regulating NE release, NPY signaling has also been reported to regulate NE biosynthesis (Wang et al., 2013). Quantitative PCR (qPCR) analyses showed that expression of *Th*, the rate-limiting enzyme in NE synthesis, was unaffected in sympathetic ganglia from NPY cko mice (**Figure 4G**), However, there was a significant up-regulation of TH expression in adrenal glands from NPY cKO mice, as revealed by qPCR and immunostaining (**Figures 4G, H**). These results are consistent with previous studies using global deletion of NPY or its receptor Y1 or pharmacological Y1R inhibition, where NPY was found to decrease TH expression in adrenal glands (Cavadas et al., 2006; Wang *et al*., 2013).

Together, these results suggest that under basal conditions, NPY derived from sympathetic neurons and/or adrenal glands, is a negative regulator of circulating NE levels, likely through the reported presynaptic inhibition of NE release, and also by blunting NE synthesis in adrenal glands.

### Loss of sympathetic NPY elicits pronounced metabolic and cardiovascular defects in mice

To address the functional consequences of loss of sympathetic-derived NPY, we performed a battery of physiological analyses in NPY cKO mice and control litter-mates. In contrast to the effects of manipulating central NPY signaling that alter body weight and food intake (Loh *et al*., 2015; Stanley and Leibowitz, 1984; Zarjevski *et al*., 1993), selective deletion of NPY from peripheral sympathetic neurons had no effect on these parameters when assessed in 1 month-old mice (**Figures 5A, B**).

**Figure 5.**
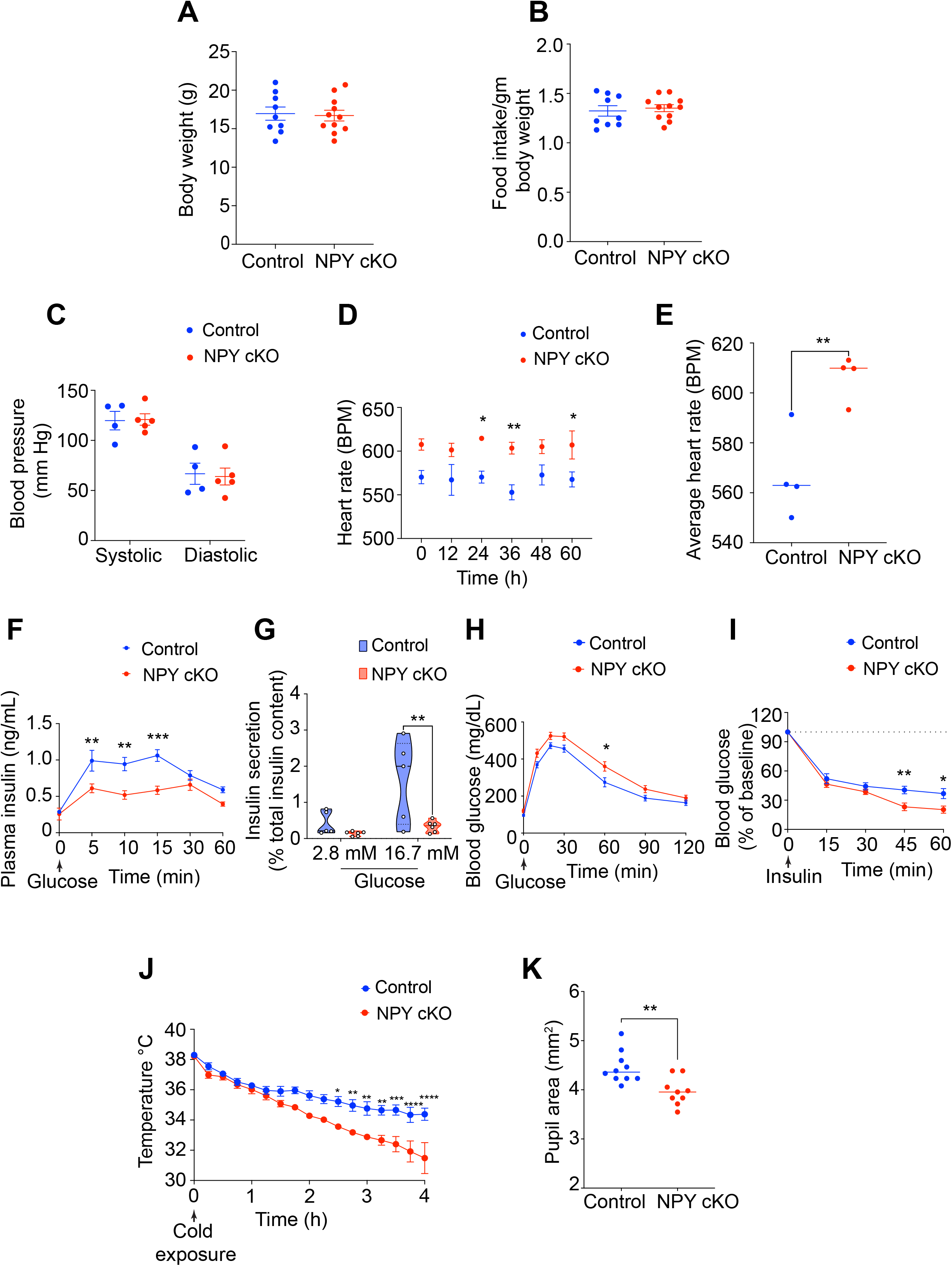
NPY cKO mice show metabolic and cardiovascular defects. **(A, B)** Body weight (A) and food intake (B) are comparable between NPY cKO and control mice. Results are means ± s.e.m from n=9 control and 11 mutant mice, unpaired t-test. **(C)** Systolic and diastolic blood pressures are normal in NPY cKO mice. Results are means ± s.e.m from n= 4 control, and 5 mutant mice, unpaired t-test. **(D, E)** NPY cKO mice show elevated heart rate, compared to control animals. ECGs were recorded continuously in conscious mice for 7 days, although only data for 4^th^-7^th^ days after insertion of lead implants are included in the analysis. Results are presented as means ± s.e.m from n= 4 mice per genotype. *p< 0.05; **p<0.01, two-way ANOVA, Sidak’s multiple comparisons tests for (D) and unpaired t-test for (E). **(F)** NPY cKO mice show reduced plasma insulin levels in response to a glucose challenge (i.p. injection of 3g/kg glucose after 16 hr fast) compared to control mice. Results are means ± s.e.m from n=12 control, 8 mutant mice. **p<0.01; ***p<0.001; two-way ANOVA, Sidak’s multiple comparisons tests. **(G)** Glucose-stimulated insulin secretion is reduced in isolated islets from NPY cKO mice relative to control animals. Islets were isolated from n= 5 control, n=6 mutant mice, p**<0.01; two-way ANOVA, Tukey’s multiple comparisons tests. **(H)** NPY cKO mice show impaired glucose tolerance. Results are means ± s.e.m from n=12 control and 17 mutant mice. *p<0.05; two-way ANOVA, Sidak’s multiple comparisons tests. **(I)** Improved insulin sensitivity in NPY cKO mice compared to control animals. Data are means ± s.e.m from n=7 control, 6 mutant. *p<0.05; **p<0.01; two-way ANOVA, Sidak’s multiple comparisons tests. **(J)** NPY cKO mice show impaired cold tolerance compared to control animals. Body temperatures were first recorded at room temperature (∼26°C), after which mice were moved to 4°C and body temperatures were measured every 15 min for 4 hr. Results are mean ± s.e.m from n=5 mice per genotype. *p<0.05; **p<0.01; ***p<0.001; ****p<0.0001; two-way ANOVA, Sidak’s multiple comparisons tests. **(K)** Dark-adapted NPY cKO mice have decreased basal pupil area compared to control mice. n=10 control, n=9 mutant mice. **p<0.01; unpaired t-test.

Blood pressure is a function of vascular resistance and cardiac output, that are both controlled by the sympathetic nervous system (Guyenet, 2006). Since NPY has been reported to be a potent vasoconstrictor, either acting directly (Ekblad *et al*., 1984; Lundberg *et al*., 1982) or potentiating the effects of NE (Dahlof *et al*., 1985; Hakanson *et al*., 1986; Lundberg *et al*., 1985), we expected that loss of sympathetic NPY would result in a drop in blood pressure. Further, NPY administration in humans (Clarke et al., 1987) or its over-expression in animals results in a long-lasting increase in blood pressure (Michalkiewicz et al., 2001). However, we found that resting systolic and diastolic blood pressures were not changed in NPY cKO mice when measuring arterial blood pressure in resting, unrestrained, and conscious mice (**Figure 5C**). In contrast to normal blood pressure, we observed a striking increase in heart rate in NPY cKO mice compared to control litter-mates, as assessed by electrocardiogram (ECG) recordings (**Figures 5D, E**). Thus, increased cardiac output might serve as a counter-acting adaptive mechanism to mask any drop in blood pressure in the absence of the vasoconstrictor effects of sympathetic-derived NPY. The sympathetic nervous system is a key contributor to the control of glucose homeostasis (Ahren, 2000; Lin et al., 2021). Sympathetic nerve-derived NE signaling rapidly elevates blood glucose levels, necessary in “fight or flight” situations, by decreasing pancreatic insulin secretion and peripheral insulin sensitivity, and by increasing pancreatic glucagon secretion (Lin *et al*., 2021). NPY administration to isolated pancreas/islets inhibits insulin secretion (Skoglund et al., 1993), while global NPY knockout mice show elevated insulin secretion (Imai et al., 2007), suggesting that it acts to potentiate NE effects. To address the role of sympathetic-derived NPY in insulin secretion, we assessed glucose-stimulated insulin secretion in NPY cKO and control mice *in vivo*. Unexpectedly, NPY cKO mice showed a pronounced reduction in plasma insulin levels in response to a glucose challenge compared to control animals (**Figure 5F**). To address whether the altered insulin secretion in NPY cKO mice was an islet-intrinsic defect, we isolated islets from control and NPY cKO mice and measured basal insulin secretion (in response to 2.8 mM glucose) and induced by high glucose (16.7 mM). We found that glucose-stimulated insulin secretion was completely abolished in NPY cKO islets (**Figure 5G**), although basal insulin secretion and total insulin content were not affected (**Figures 5G and Figure S2A**). The observations in isolated islets from NPY cKO mice suggest that loss of sympathetic nerve-derived NPY results in islet-intrinsic changes that then attenuate insulin secretion. Of note, NPY expression is downregulated in mature islets (Rodnoi *et al*., 2017; Teitelman et al., 1993), and DBH is not expressed in islet cells (see **Figure S1A**), suggesting that the insulin secretion defect in NPY cKO islets is not due to NPY loss in islet cell types. Consistent with the blunted insulin secretion in NPY cKO mice, we found that mutant mice had impaired glucose tolerance compared to control animals (**Figure 5H**). However, the glucose intolerance was milder than expected from the striking decrease in glucose-stimulated insulin secretion *in vivo* and *in vitro*. One potential explanation is that we found that NPY cKO mice showed improved insulin sensitivity compared to control litter-mates (**Figure 5I**). Together, these results reveal that sympathetic-derived NPY controls glucose homeostasis, in part, by positively regulating islet insulin secretion. Further, the improved insulin sensitivity in NPY cKO animals may reflect a compensatory response to the low plasma insulin levels and/or a decrease in noradrenergic signaling in insulin-responsive peripheral organs, including liver, adipose tissue, and striated muscles in the absence of sympathetic NPY. The sympathetic nervous system is a critical mediator of physiological responses to cold exposure through triggering piloerection, vasoconstriction, and non-shivering thermogenesis in the brown adipose tissue (Tan and Knight, 2018). To address if NPY expressed in sympathetic neurons is required to maintain body temperature in response to cold exposure, we performed a cold tolerance test in NPY cKO and control animals. Mice were implanted with sub-cutaneous temperature sensors, and body temperatures were first recorded at room temperature, following which mice were moved to 4°C and body temperatures measured every 15 min for 4 hr. While both NPY cKO and control animals were able to maintain their internal temperatures for the first hr at 4°C, the body temperatures in mutant mice dropped sharply compared to the controls with prolonged cold exposure (**Figure 5J**). Thus, sympathetic-derived NPY is required for animals to mount appropriate physiological responses to cold. Together, with our observations that NPY cKO animals fail to show an increment in circulating NE in response to a brief cold stimulus (4°C, 2 hr) (**Figure 4F**), these results suggest that stress-induced responses are impaired in mutant animals. Lastly, we assessed pupil size, which is controlled by a balance between the pupil dilator muscle, innervated by sympathetic fibers from the SCG, and the pupillary sphincter muscle, innervated by parasympathetic nerves coming from the ciliary ganglion (McDougal and Gamlin, 2015). Sympathetic nerve-derived NE signaling drives the contraction of smooth muscle cells in dilator muscle (McDougal and Gamlin, 2015). To measure basal pupil size, control and NPY cKO mice were dark-adapted for over 2 hr, and pupil sizes were recorded for 10 seconds in the dark under non-anesthetized conditions. We observed decreased basal pupil areas in NPY cKO mice compared to controls (**Figures 5K and Figure S2B**). To ask whether this phenotype is due to decreased parasympathetic activity, we measured pupil constriction in response to increasing light intensities, ranging from 0.01-1000 lux, administered for 30 seconds. Light onset at 0.1 lux or higher resulted in rapid constriction with greater constrictions at higher light intensities in both NPY cKO and control mice (**Figure S2C**). The intensity responses were virtually identical for the two groups (**Figure S2C**). These results suggest that parasympathetic function is intact in NPY cKO mice, and that the decreased pupil areas likely reflect reduced noradrenergic signaling at the pupil dilator muscle with the loss of sympathetic-derived NPY.

## Discussion

Despite decades of research on NPY, specifically for its actions in the brain, the peripherally-mediated functions of NPY have remained relatively under-appreciated. Here, using conditional knockout mice, we reveal specific roles for NPY expressed in peripheral sympathetic neurons in controlling neuron soma size during development and in regulating autonomic physiology, including cardiovascular function, blood glucose homeostasis, cold tolerance, and pupil size, in adult animals.

In contrast to the canonical view that NPY primarily functions as a modulator of sympathetic neurotransmission (Lundberg et al., 1989b; Zukowska-Grojec, 1998), our results suggest a more complex scenario. For example, NPY cKO mice showed decreased glucose-stimulated insulin secretion and impaired glucose tolerance (**Figures 5F-H**), which are contrary to that predicted from the loss of NPY-mediated potentiation of NE signaling, where NE is known to potently inhibit insulin secretion and elevate blood glucose (Ahren, 2000). Strikingly, the phenotypes in NPY cKO mice recapitulate the effects that we previously observed with developmental ablation of sympathetic nerves (Borden et al., 2013), which resulted in reduced insulin secretion and glucose intolerance as well as disruptions in islet architecture in mice (Borden et al., 2013). Since NPY is removed early during embryonic development in NPY cKO mice, these findings suggest that nerve-derived NPY might exert a developmental role in regulating islet morphology and function. Unexpectedly, we also observed an increase in heart rate in NPY cKO animals, which at first glance, was incongruent with a canonical function for NPY in potentiating NE effects. However, given the normal blood pressure in NPY cKO mice, we reason that the tachycardia phenotype may be due to a compensatory cardiac self-regulatory response aimed to maintain the blood pressure constant in a condition of low vascular resistance when NPY is removed from sympathetic nerves. Alternatively, the elevated heart rate in mutant animals could reflect a direct role for sympathetic-derived NPY in reducing heart rate through inhibition of parasympathetic cholinergic signaling (Herring et al., 2008). Other phenotypes observed in NPY cKO mice, for example, the improved insulin insensitivity, impaired cold tolerance, and reduced basal pupil size, are all consistent with decreased sympathetic tone and the reported canonical role of NPY in potentiating NE signaling. Together, our results suggest that NPY derived from sympathetic neurons, can act alone or in concert with the classical neurotransmitter, NE, in a target tissue-dependent context.

Of note, NPY and NE are co-expressed in the nucleus of the solitary tract (NTS) (Chen et al., 2020; Everitt et al., 1984) and A1/C1 cluster of catecholaminergic neurons in the brainstem (Liu *et al*., 2020; Sawchenko et al., 1985), which have been implicated in regulating food intake (Chen *et al*., 2020; Sahu et al., 1988) and sympathetic outflow to the periphery (Tseng et al., 1989; Vahatalo et al., 2015). While we cannot exclude the possibility that loss of NPY in these brain nuclei may account for some of the phenotypes that we observed in NPY cKO mice, our results suggest that pre-ganglionic input to sympathetic ganglia neurons was unaffected based on c-FOS immunostaining (**Figures 4A-D**) and expression of catecholamine synthetic enzymes such as TH (**Figure 4G**). Further, food intake was normal in mutant animals. Further studies are warranted to fully dissect the roles of NPY and NE co-transmission in the brain versus periphery. Axonal terminations of NPY-expressing sympathetic neurons contain small NE-containing vesicles, which are rapidly released upon acute stimulation, as well as large dense core vesicles containing both NE and NPY that slowly release both molecules under prolonged or high frequency stimulation (De Potter et al., 1997; Lundberg *et al*., 1989a). The findings that NPY is specifically released during intense and prolonged high-frequency activation has led to the view that NPY is poised to regulate sympathetic responses in extreme physiological circumstances, including chronic stress, exposure to danger, and pathological conditions such as heart failure and insulin-induced hypoglycemia (Hirsch and Zukowska, 2012; Michalkiewicz *et al*., 2001; Tan *et al*., 2018). Yet, we found that loss of NPY from sympathetic neurons results in pronounced autonomic defects, including increased heart rate, impaired insulin secretion, glucose intolerance, and reduced pupil size, under basal conditions. These results reveal that, even under basal conditions, sympathetic-derived NPY exerts a wide range of physiological effects across a number of organ systems. It is possible under basal conditions, an increase in circulating NE in the absence of sympathetic-derived NPY contributes to some of the phenotypes that we observed in the mutant mice. Together, with previous studies, our results suggest that co-existence of NPY with NE in sympathetic nerve terminals and adrenal glands may serve as a buffer to protect tissues from the adverse effects of excessive or chronic NE release.

The NPY system has been widely recognized as one of the most important regulators of whole-body physiology in animals. Yet, despite decades of study, dissecting the tissue- and cell-specific roles of NPY *in vivo* through the use of genetic tools has lagged behind. Here, through selective deletion of NPY from peripheral sympathetic neurons, we provide direct evidence that sympathetic-derived NPY is essential for metabolic and cardiovascular regulation in mice. In humans, increased circulating NPY has been documented in several pathological conditions where sympathetic output is also increased, including hypertension, chronic heart failure, and insulin resistance (Jaakkola et al., 2010; Solt et al., 1990; Tan *et al*., 2018). Our findings suggest that targeting peripheral NPY signaling with specific agonists or antagonists can offer possibilities in the treatment of metabolic and cardiovascular diseases.

## Acknowledgements

We thank Haiqing Zhao for helpful comments on the manuscript. We also thank Ivo Kalajzic (UConn Health) for the generous gift of the *NPY*^fl/fl^ mice. We thank the JHU Integrated Imaging Center for assistance with microscopy. This work was supported by NIH R01 awards, NS114478 and NS107342, to R.K.

## Author Contributions

R.P., R.K., and R.K. designed the study. R.P. and R.K. conducted the majority of the experiments and analyzed data. H.W., M.Y. and S.H., provided guidance with pupil analyses and temperature recordings. K.Q. assisted with whole-organ immunohistochemistry. S.B. and E.T. performed the heart rate and blood pressure analyses. R.P., R.K, and R.K. wrote the manuscript.

## Supplemental Figure Legends

**Figure S1.**
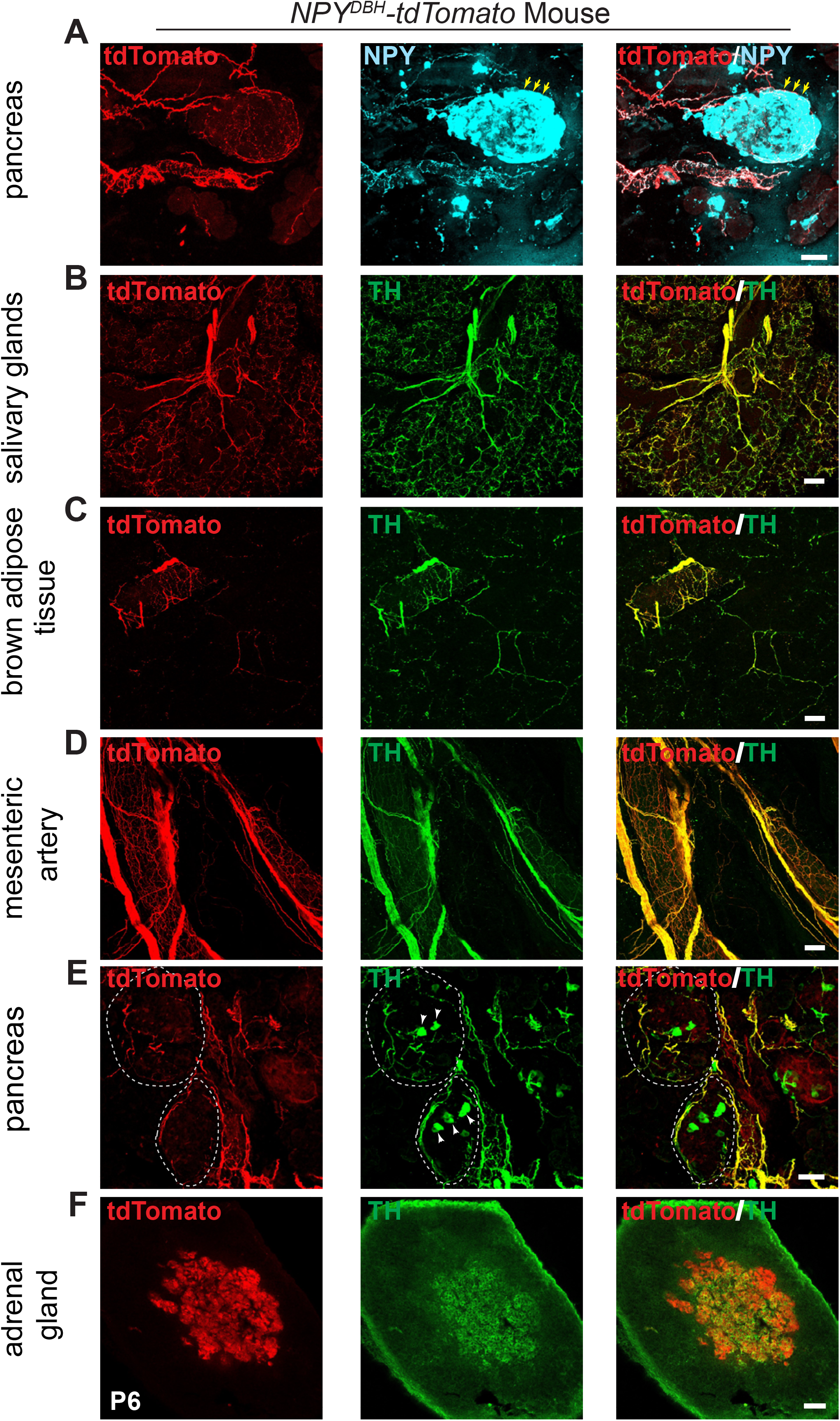
Innervation of peripheral organs by NPY^DBH^-tdTomato sympathetic axons. Related to Figure 2. **(A)** TdTomato reporter signal (red) is observed in axons innervating pancreatic islets, but not seen in islet endocrine cells which show intrinsic NPY immunoreactivity (cyan, yellow arrows) in NPY^DBH^-tdTomato mice. (**B, C**) Partial overlap of TH immunoreactivity with tdTomato reporter signal in axons innervating paravertebral targets, salivary glands (B) and BAT (C). **(D-F)** Complete overlap of tdTomato reporter expression with TH immunostaining in mesenteric artery (D) and pancreatic islets (E), which are prevertebral targets, as well as adrenal chromaffin cells (F). Note that TH is also expressed in some islet endocrine cells (white arrowheads in E). Scale bars, 50 μm.

**Figure S2.**
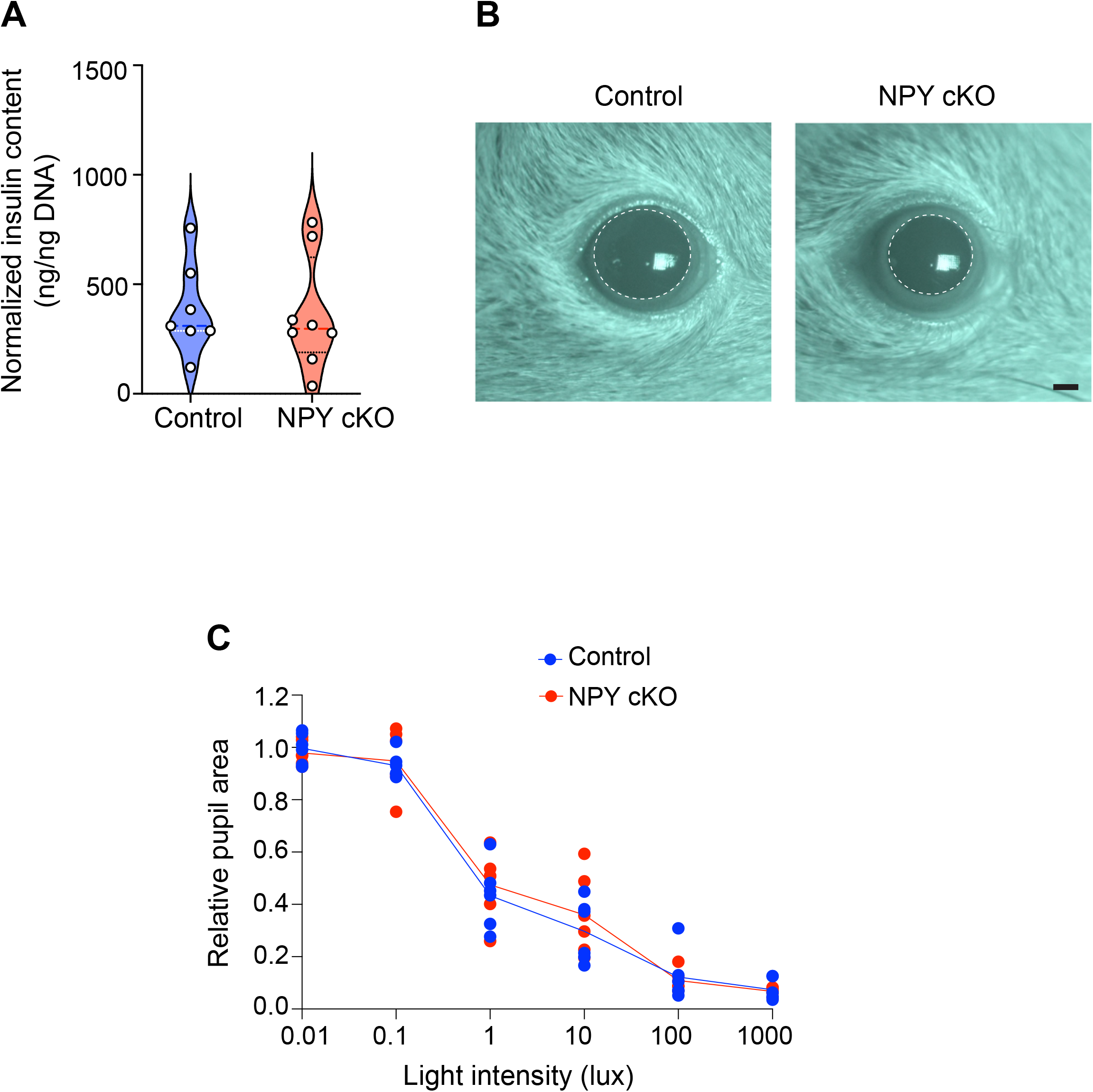
Additional functional analyses in NPY cKO mice. Related to Figure 5. **(A)** Total insulin content in pancreatic islets is unaffected in NPY cKO mice. Insulin content is normalized to total islet DNA content. Results are means ± s.e.m from islets isolated from n= 7 control and 8 mutant mice, unpaired t-test. **(B)** Representative pupil images of control and mutant mice. Pupil areas are shown by white dotted circles. Mutant mice show reduced pupil size. Scale bar, 160 μm. **(C)** Pupil constriction in response to increasing light intensities is comparable between control and mutant mice indicating normal parasympathetic activity in NPY cKO mice. Results are means ± s.e.m. from n=6 mice per genotype, two-way ANOVA with Bonferroni’s correction.

## STAR METHODS

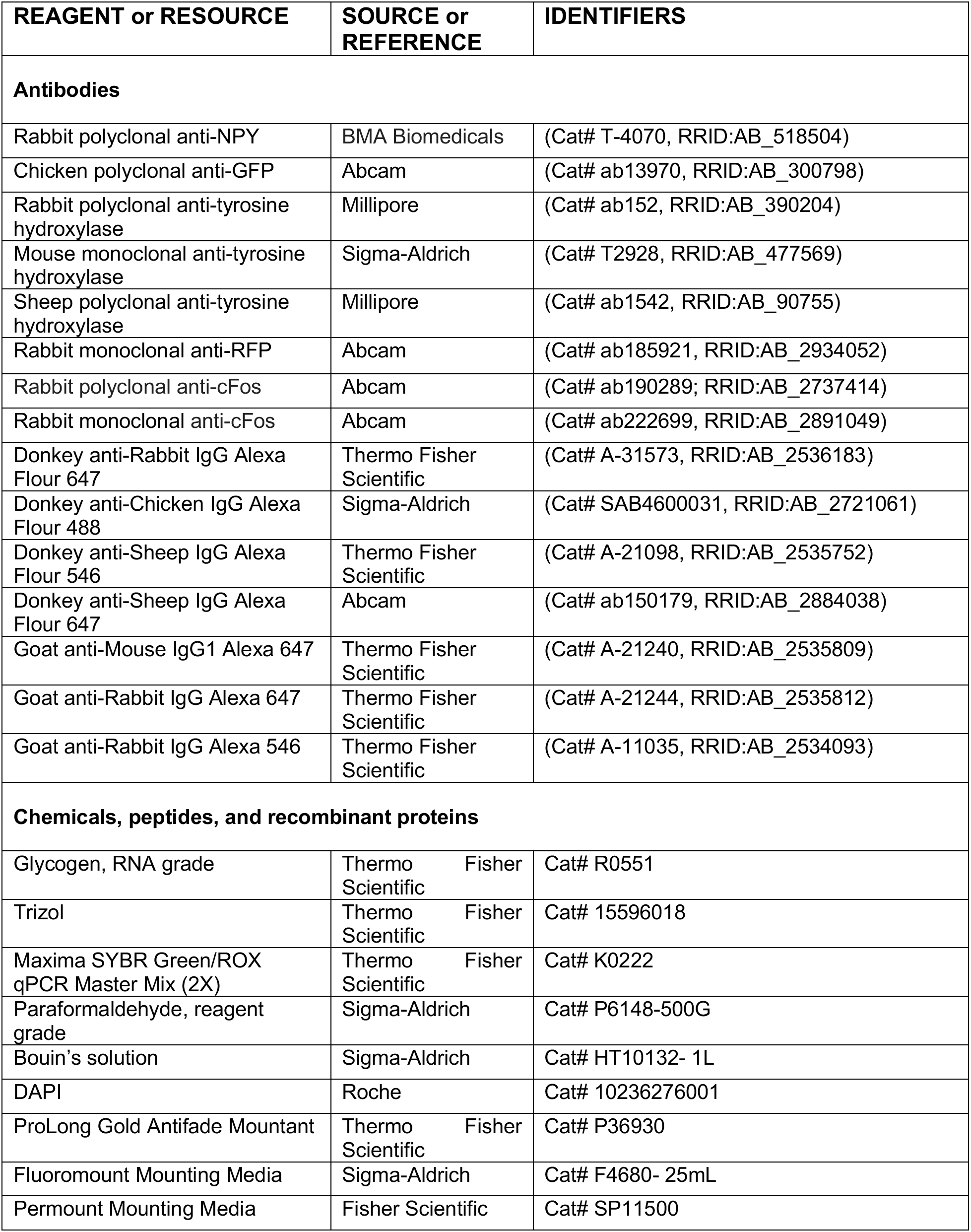
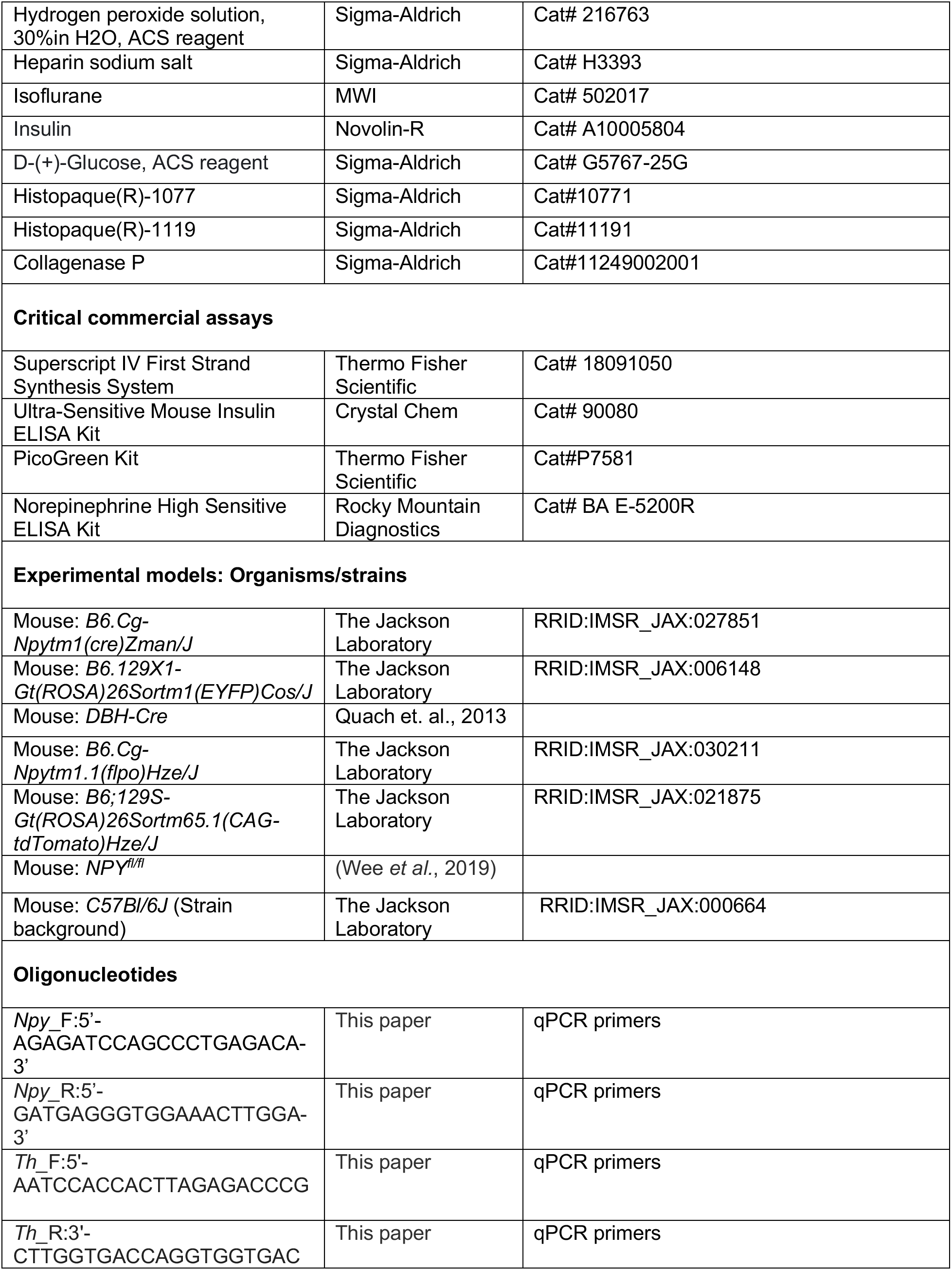
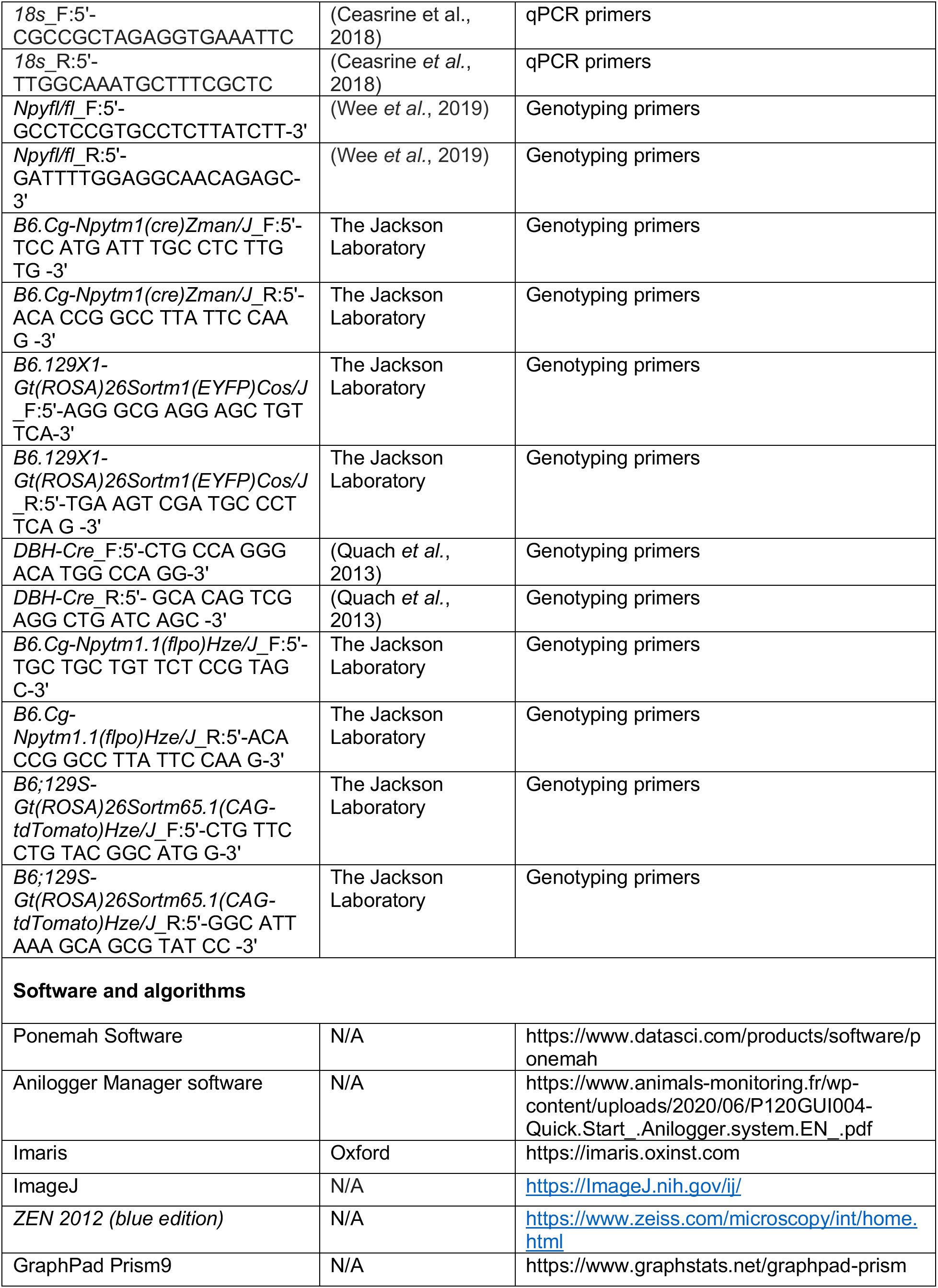

### RESOURCE AVAILABILITY

#### Lead Contact

Further information and requests for resources and reagents should be directed to and will be fulfilled by the Lead Contact, Rejji Kuruvilla.

#### Materials Availability

All transgenic mice generated in this study (see **Key Resources**) are available upon request.

#### Data and Code Availability

This paper does not report original code.

Any additional information required to re-analyze the data reported in this paper is available from the lead contact upon request

### EXPERIMENTAL MODEL AND SUBJECT DETAILS

#### Animals

All animal care and experimental procedures were conducted in accordance with the Johns Hopkins University Animal Care and Use Committee (ACUC) and NIH guidelines. All efforts were made to minimize the pain and number of animals used. Animals were group housed in a standard 12:12 light-dark cycle, except for pupil analysis where the animals were dark-adapted for two days, with excess water and food ad libitum. The ages of mice are indicated in the figure legends and/or methods. Both male and female mice were used for the analysis. The following lines were used in this study: *NPY^Cre^ (B6.Cg-Npytm1(cre)Zman/J,* JAX #027851), *ROSA26^eYFP^ (B6.129X1-Gt(ROSA)26Sortm1(EYFP)Cos/J,* JAX #006148), *NPY-Flp (B6.Cg-Npytm1.1(flpo)Hze/J,* JAX #030211), and *ROSA26^tdTomato^ (B6;129S-Gt(ROSA)26Sortm65.1(CAG-tdTomato)Hze/J* JAX #021875) mice were all obtained from The Jackson Laboratory. *DBH^Cre^* and *NPY*^fl/fl^ mice were generous gifts from Dr Warren G. Tourtellotte (Northwestern University) and Dr. Ivo Kalajzic (UConn Health), respectively.

## Method Details

### Isolation of RNA and quantitative RT-PCR analysis

Total RNA was isolated from dissected SCG, CG-SMG, and adrenal glands using the TRIzol-chloroform extraction method. cDNA was prepared using Superscript IV First-Strand Synthesis System. Real-time qPCR analysis was performed using a Maxima SYBR green/Rox Q-PCR Master Mix (Thermo Fisher Scientific) and gene-specific primers, in a 7300 real-time PCR system (Applied Biosystem). All samples were analyzed in triplicate reactions. Fold change was calculated using the 2 ^(ΔΔCt)^ normalized to 18S transcript. Primer sequences are listed in the **Key Resource** table.

### Food intake

Mice (4-5 weeks) were weighed and then individually housed for a total of 7 days. 100gm of food ad libitum was given to each mouse on the first day. The uneaten food was measured each day, total food intake was calculated and normalized to the body weight per mouse.

### Norepinephrine (NE) ELISA

Blood samples (300 μl) were drawn retro-orbitally from anesthetized mice (6 weeks) that were either housed at room temperature or exposed to cold (4°C, 2 hr). Plasma was prepared by centrifuging blood at 3000 rpm for 15 min, 4 °C, and 100 μl plasma was used for NE measurements using ELISA (NE High Sensitive ELISA Kit, Rocky Mountain Diagnostics) according to manufacturer’s protocol.

### Blood pressure recordings

Blood pressure was measured by a non-invasive system (BP-2000 Series II, Visitech systems). Mice (6-8 weeks) were acclimatized to the machine for two days prior to blood pressure recordings to minimize stress. Each mouse was placed inside a magnetic restrainer and an inflatable occlusion cuff with a light sensor that was passed through the tail. Systolic and diastolic blood pressures were calculated based on the variations in the amount of light transmitted through the tail. 20 blood pressure recordings were averaged to obtain the final measurements for each mouse.

### Electrocardiograms

ECG recordings were performed on adult mice as previously described (Mapps et al., 2022). Briefly, mice (6-8 weeks) were anesthetized with 4% isoflurane, intubated, and placed on ventilator support (settings 1.2 ml/g/min at 80 breaths/min). The animal’s abdomen was shaved, scrubbed with betadine and alcohol, and draped with a sterile barrier with the surgery site exposed. A small 0.5 cm midline incision was performed and a telemetry unit (DSI Harvard Bioscience Inc.) was implanted intraperitoneally. ECG leads were implanted subcutaneously and sutured over the right upper and left lower chest. Body temperature was maintained at 37°C. Immediately following implantation, the wound was closed with a 3-0 silk suture. Anesthesia was turned off and the animal was monitored for spontaneous breathing and was given a subcutaneous buprenorphine injection (0.01–0.05 mg/kg buprenorphine, IM) to alleviate pain. ECGs were subsequently recorded continuously in conscious animals for approximately 7 days using the Ponemah Software (DSI Harvard Bioscience Inc.). Mice were kept at a stable temperature with a regular 12 h light/dark cycle. To exclude the effects of pain and anesthesia, continuous ECG recordings between days 4 and 7 post-lead implantations were only included in the analysis of mean heart rates.

### Cold tolerance

Mice were acclimatized for at least 3 weeks at 26°C. Mice at 6-8 weeks of age were anesthetized with isoflurane, abdominal skin was shaved, and a 1.5 cm incision was made at the abdominal midline. Real-time readable temperature loggers (Anipill, Herouville, France) were implanted into the intraperitoneal cavity to a position ventral to the caudal arteries and veins but dorsal to the abdominal viscera. Abdominal musculature was closed with an absorbable suture (chromic gut 4/0; AD surgical) and the skin incision was with a non-absorbable nylon (Ethilon black 4/0; ProNorth Medical) suture. Analgesia (meloxicam 2mg/Kg) was provided subcutaneously for 2 days following surgery. After recovery for 10-12 days from surgery, mice were transferred to a 4°C cold cabinet for 4 hr, where they were individually housed. Core body temperatures were recorded every 15 minutes using the AniLogger Monitor. Data were analyzed using GraphPad Prism 9.

### Glucose-induced insulin secretion

Mice (6-8 weeks) were individually housed and fasted overnight (16 hr). The next morning, mice were i.p. injected with 3 g/kg glucose/saline (Sigma-Aldrich). Tail blood was collected at the indicated time, spun down at 3000 rpm for 15 min, and plasma insulin levels were measured with an Ultrasensitive Insulin ELISA kit (Crystal Chem, Elk Grove Village, IL).

### Mouse islet isolations, in vitro insulin secretion and insulin content

Islets were isolated from mice at 6-8 weeks of age as previously described (Wollheim et al., 1990). Briefly, islets were isolated by collagenase distension through the bile duct (Collagenase P [Roche, Basel, Switzerland], 0.375 mg/mL in HBSS) and digestion at 37°C. Digested pancreata were washed with HBSS +0.1% BSA and subjected to discontinuous density gradient using histopaque (6:5 Histopaque 1119:1077; Sigma, St. Louis, MO). The islet layer (found at the interface) was collected and islets were handpicked under an inverted microscope for subsequent analysis. For insulin secretion in isolated islets, harvested islets were allowed to recover overnight in RPMI-1640 media containing 10% fetal bovine serum, and 5U/L penicillin/streptomycin (Invitrogen). Islets were washed with Krebs-Ringer HEPES buffer (KRBH) containing low (2.8 mM) glucose and pre-incubated for 1 h in low-glucose KRBH. After pre-incubation, groups of similarly sized 14 islets were handpicked into a 24-well plate and allowed to incubate for 30 min in KRBH containing low glucose, or high (16.7 mM) glucose. After incubation the supernatant fractions were removed and the islets were lysed with acid ethanol, followed by ELISA to determine the insulin content in both supernatant and islet fractions. Insulin values were normalized to islet DNA content from the same lysates using a Picogreen Kit (Thermo Fisher Scientific).

### Glucose tolerance test

Mice (6-8 weeks) were individually housed and fasted overnight (16 hr). The next morning, mice were i.p. injected with 2g/kg glucose/saline (Sigma-Aldrich), and tail blood glucose levels were measured using OneTouch Ultra glucometer at the indicated time.

### Insulin sensitivity test

Mice (6-8 weeks) were individually housed and fasted for 6 h before being i.p. injected 0.75 U/kg of insulin (Novolin-R, Novo Nordisk, Princeton, NJ). Tail blood glucose levels were measured using a OneTouch Ultra glucometer at indicated times as previously described (Ceasrine *et al*., 2018).

### Pupil analyses

Pupil size measurements were performed as reported previously (Keenan et al., 2016). Mice were dark-adapted for over 2 hours before the experiment. For all recordings, mice were un-anesthetized and restrained by hand. To mitigate stress, which can affect pupil size, researchers handled mice for several days prior to the experiments. A Sony Handycam (FDR-AX33), carrying a prime lens, set at manual focusing and night shot mode, accompanied by an external infrared light source, was used for recording. The dotted pattern from the infrared light source is reflected by the mouse cornea and used as the focusing indicator. The equipment ensures that mouse pupils can only be focused at a fixed distance from the camera and can be easily visualized in the dark. Pupil size was measured by the maximum diameter using Fiji software in each picture frame. To measure dark-adapted baseline pupil size, a 10 second video was recorded.

To examine parasympathetic activity, mice were dark-adapted as described above. Un-anesthetized mice were restrained by hand. To measure pupil responses to light, a video of 10 sec in the dark followed by 30 sec exposure to white light at 0.01, 0.1, 1, 10, 100, 1000 lux, respectively, were recorded. The pupil constriction is quantified by normalizing the pupil area at 30 second to the baseline area in the dark.

### Neuronal counts

Neuronal counts were performed as previously described (Mapps *et al*., 2022). In brief, CG-SMGs were dissected from mice at P21, fixed in 4% PFA at 4°C for 4 hr, and cryoprotected in 30% sucrose/PBS overnight. CG-SMGs were mounted in OCT and serially sectioned with 12 μm thickness. Every fifth section was stained with a solution containing 0.5% cresyl violet (Nissl). Cells in CG-SMGs with characteristic neuronal morphology and visible nucleoli were counted using ImageJ.

### Immunohistochemical analyses

For SCG, and CG-SMG immunostaining, mice torsos at indicated ages were fixed in 4% PFA/PBS for 4 hr to overnight and cryoprotected in 30% sucrose/PBS for 2-3 days. Target tissues from mice at postnatal day 6 or 21, were dissected and fixed in 4% PFA/PBS overnight and cryoprotected in 30% sucrose/PBS for 48 h. Torsos and target tissues were embedded in OCT (Sakura Finetek) and stored at −80°C. SCG, CG-SMG were cryo-sectioned at 14 μm while target tissues at 50 μm. For paraffin embedding, SCG and CG-SMG were fixed in Bouin’s solution for 1 hr at room temperature, washed, and left in 70% ethanol at room temperature, followed by dehydration with 50%, 75%, 85%, 95%, and 100% ethanol and xylene. Tissues were embedded in paraffin and sectioned at 6 μm using a microtome, deparaffinized using xylene, and rehydrated using 100%, 95%, 85%, 75%, and 50% ethanol. Paraffin sections were incubated in 10 mM sodium citrate buffer (pH 6) at 95°C for 10 min, followed by incubation in 0.2 M glycine for 10 min. Both cryo- and paraffin sections were then permeabilized in 1% Triton X-100 in PBS for 10 min at room temperature. Blocking was done in 10% goat or donkey serum/3% bovine serum albumin in 0.3% Triton-X 100 in PBS for 1-2 h at room temperature. Primary antibodies used were rabbit/sheep/mouse anti-TH (1:300), rabbit anti-NPY (1:250), chicken anti-GFP (1:500), rabbit anti-RFP (1:500), with incubations overnight at 4 °C. Slides were washed with 0.3% Triton-X 100/PBS and incubated with Alexa-488, −546, or −647 conjugated anti-rabbit, anti-sheep, anti-mouse or anti-chicken secondary antibodies (1:400-500) in blocking solution for 1-2 hr at room temperature. Tissues were then mounted in Aqueous Mounting Medium containing 100 μg/ml DAPI and imaged at 1 μm optical slices using a Zeiss LSM 700 or LSM 800 confocal scanning microscope. Maximum-intensity images were generated using ImageJ.

For cFos immunostaining, CG-SMG was dissected from mice that were kept at room temperature or exposed to cold (4°C, 1 hr). Tissues were cryoprotected in 30% sucrose/PBS for 1hr at 4 °C and embedded in OCT and stored at −80°C. 16 μm thick sections were prepared and fixed with 2% PFA/PBS for 5 min and permeabilized with freshly prepared 1% Triton-X-100 for 15 min. Blocking was done in 5% goat serum/1% bovine serum albumin in 0.3% Triton-X100/PBS for 1.5 h at room temperature, followed by incubation with a polyclonal or monoclonal anti-rabbit cFos (1:1000) antibody diluted in blocking solution overnight at 4 °C. After washes, sections were incubated in Alexa-546 conjugated anti-rabbit secondary antibody overnight at 4 °C. Sections were mounted using ProLong Gold (Thermo Fisher) antifade mounting media and images were acquired using LSM 800 confocal scanning microscope.

### iDISCO and wholemount immunostaining

iDISCO-based tissue clearing for whole mount immunostaining of sympathetic chain and target organs from P6 mice was performed as previously described (Scott-Solomon and Kuruvilla, 2020). Briefly, the sympathetic chain and organs were fixed in 4%PFA/PBS, dehydrated by methanol/water series (20-80%), and incubated overnight in 66% dichloromethane (DCM)/33% methanol. Tissues were bleached with 5% H_2_O_2_ in methanol at 4°C overnight, re-hydrated with methanol/PBS/0.2% Triton-X100 series (80-20%), and permeabilized first with 0.2%TritonX-100/PBS for 2 h followed by 3 overnight permeabilizations with 0.2% TritonX-100/20%DMSO/0.3M glycine in PBS. Tissues were incubated in blocking solution (0.2% TritonX-100/10% DMSO/6% Normal Goat/Sheep Serum in PBS) for 2 overnight and then incubated with rabbit/sheep-anti-TH (1:300) or rabbit anti-RFP (1:300) in 0.2% Tween-20/0.001% heparin/5% DMSO/3% Normal Goat/Sheep Serum in PBS at 37 °C for 4 overnight. Samples were then washed with 0.2% Tween-20/0.001% heparin in PBS and incubated with anti-rabbit Alexa-546 or anti-sheep Alexa-647 secondary antibody (1:400) in 0.2% Tween-20/0.001% heparin/3% Normal Goat/Sheep Serum in PBS for 4 overnight. Tissues were extensively washed with 0.2% Tween-20/0.001% heparin in PBS and dehydrated in methanol. Tissues were cleared by successive washes in 66% DCM/ 33% methanol, 100% DCM and 100% Dibenzyl Ether. Tissues were imaged using a light-sheet microscope (LaVision BioTec Ultra Microscope II). Imaris software was used for 3D reconstructions of images.

For wholemount immunostaining of ganglia, SCG or CG-SMG were dissected from P6 mice and fixed with 4%PFA/PBS at 4°C for 4 hr. Tissues were permeabilized using 0.2% TritonX-100/20%DMSO/0.3M glycine/PBS for 1 hr at room temperature and blocked with 0.2% TritonX-100/10% DMSO/6% Normal Goat Serum at room temperature overnight. Tissues were then incubated with primary antibody anti-rabbit TH antibody (1:300) in 0.2% Tween-20/0.001% heparin/5% DMSO/3% Normal Goat Serum at room temperature, 2 overnight. Tissues were washed with 0.2% Tween-20/0.001% heparin/PBS extensively and incubated with anti-rabbit Alexa-647 secondary antibody (1:400) in 0.2% Tween-20/0.001% heparin/3% Normal Goat Serum/PBS at room temperature, 2 overnight. tdTomato fluorescence was visualized with endogenous expression. Tissues were mounted using Aqueous Mounting Medium and images were acquired using LSM 800 confocal microscope.

### Quantification and statistical analysis

For practical reasons, analysis of soma-size measurements, and neuron numbers analysis were performed in a semi-blinded manner. The experimenter was aware of the genotypes prior to the experiment but performed each immunostaining and analyses without knowing the genotypes. All physiological analyses were performed in a blinded manner with the experimenter only keeping track of the ear tag numbers. All graphs and statistical analyses were performed using GraphPad Prism 9. Statistical significance was determined using an unpaired, two-tailed student t-test, and a two-way ANOVA test for more than one variable. All error bars are represented as the standard error of the mean (SEM).

